# Longitudinal blood DNA methylation profiling reveals disrupted immune–epigenetic adaptation and candidate stress-related loci in postpartum depression

**DOI:** 10.64898/2026.04.03.716376

**Authors:** Philip Wolff, Elena Losse, Susanne Nehls, Geraldine Zimmer-Bensch, Natalia Chechko

## Abstract

Postpartum depression (PPD) arises during a period of profound endocrine and immune reorganisation, yet it is unclear whether women who develop PPD show distinct trajectories of immune-related DNA methylation compared to euthymic mothers. In a longitudinal cohort, women with PPD (n = 17) and healthy postpartum controls (n = 24) were followed from birth to 12 weeks postpartum, with repeated assessment of depressive symptoms and perceived stress and whole-blood sampling at 2–3 days (T0) and 12 weeks (T4) for Infinium MethylationEPIC array profiling. Healthy postpartum women showed a widespread gain in DNA methylation from T0 to T4 with strong enrichment of genes involved in neutrophil activation, chemokine signalling and interleukin-1 production, consistent with a normative immune-epigenetic down-tuning after childbirth. Women with PPD also exhibited immune-related changes, but with fewer differentially methylated CpGs and increased variance at sites that were stably hypermethylated in controls, indicating an attenuated and more heterogeneous epigenetic response. Although no CpG reached epigenome-wide significance in direct case-control contrasts, longitudinal consistency analyses highlighted a small set of CpGs with reproducible PPD-associated hypermethylation in stress- and signalling-related genes, including FKBP5 and AVP, suggesting that disrupted immune-epigenetic adaptation and altered regulation at these loci may contribute to postpartum vulnerability.

## 1. Introduction

During the early postpartum phase, particularly within the first three months of delivery, women undergo substantial biological changes across multiple physiological systems. These include rapid hormonal shifts, marked alterations in brain structure and function (1–3), adaptations of the hypothalamic-pituitary-adrenal (HPA) axis (4), and dynamic changes in immune regulation (5). These abrupt endocrine transitions occur in parallel with widespread neurobiological and systemic adaptations, which support maternal recovery and facilitate the transition to caregiving. At the same time, this period of pronounced biological plasticity may also increase vulnerability to affective disorders. Up to 10% of new mothers develop postpartum depression within the early postpartum period (6,7), a condition that is associated with impairments in maternal caregiving behavior and may have long-term consequences for child development (8–14).

Biological research increasingly highlights that pregnancy and the postpartum period represent states of profound immune recalibration, during which pregnancy-induced immune adaptations gradually shift back toward a non-pregnant state (15). These postpartum immune system changes are thought to be accompanied by dynamic epigenetic modifications (Effects of postpartum hormonal changes on the immune system and their role in recovery - PubMed; Risk Factors for Postpartum Depression Based on Genetic and Epigenetic Interactions - PubMed). Converging evidence further implicates immune and inflammatory pathways in the pathophysiology of major depression and related mood disorders, suggesting that epigenetic alterations in immune-related genes may play a critical role in maternal mental health (16).

Consistent with these observations, women with PPD frequently show elevated levels of pro-inflammatory cytokines such as IL-6, IL-1β, and TNF-α. Sustained low-grade inflammation has also been found to interfere with neural plasticity, monoaminergic signaling, and stress-related pathways, thereby promoting depressive symptoms (16–18). However, the biological mechanisms linking postpartum immune adaptation to PPD remain incompletely understood, and are unlikely to be explained by genetic factors alone. Epigenetic mechanisms, on the other hand, provide a compelling link between environmental exposures, hormonal fluctuations, and long-lasting gene regulation, with DNA methylation serving as a key molecular interface between external signals and genomic function (19).

Prior work has identified altered methylation of candidate genes such as the glucocorticoid receptor (NR3C1) and oxytocin receptor (OXTR) in association with maternal depression, suggesting broader epigenetic reprogramming in stress- and immune-related pathways in peripartum women (20–22). Yet, dynamic DNA methylation changes across the postpartum period remain poorly characterized, particularly with regard to their longitudinal trajectories in women who develop PPD compared to those who remain euthymic, as most existing studies have focused on single time points. Importantly, part of the longitudinal methylation remodeling observed in whole blood may be driven or amplified by physiological postpartum immune reconstitution, highlighting the need to account for such processes when investigating postpartum epigenetic trajectories (5).

To address this, the present study investigated women with PPD and healthy postpartum controls longitudinally at two postpartum time points, enabling the characterization of methylation trajectories while accounting for physiological postpartum adaptation. Blood samples were obtained shortly after childbirth (2-3 days, T0) and at 12 weeks postpartum (T4), allowing us to capture epigenetic remodeling during a critical window of immune and endocrine readjustment and compare the coordination and variability of methylation trajectories between the groups.

We hypothesized that (i) women who develop PPD would show distinct longitudinal methylation patterns in stress- and immune-related genes compared to healthy postpartum women, and (ii) these differences would reflect not only persistent epigenetic alterations associated with depressive symptoms but also interactions with physiological postpartum immune reconstitution. The study therefore focuses on identifying disorder-specific methylation trajectories while disentangling them from normative postpartum epigenetic changes.

## 2. Materials / Subjects and Methods Cohorts and sample description

Between 2018 and 2023, N = 827 euthymic women (age range 23 to 43) were recruited within 6 days of delivery with written informed consent (t0). Following inclusion, the participants’ sociodemographic information, information surrounding the pregnancy and childbirth, medical history and personal and family psychiatric history were obtained. A blood sample was drawn to analyze DNA methylation profiles. Subsequently, the participants were contacted at three-week intervals (t1 to t4) to assess their mood (Edinburgh Postnatal Depression Scale; EPDS), stress level (Perceived Stress Scale; PSS-10), and attachment to the infant (Maternal Postnatal Attachment Scale; MPAS) over a period of 12 postpartum weeks. In addition, using the online software “Survey Monkey”, the subjectively perceived stress level was queried every two days on a ten-point Likert scale (1 = “no subjective stress” to 10 = “very high subjective stress”) to facilitate a continuous mapping of the individual stress experience over the entire period of observation.

At 12 weeks postpartum (t4), a second blood sample was drawn and a clinical interview was performed to assess the participant’s mood and the likelihood of a diagnosis of postpartum depression (PPD).

A complete blood sampling protocol was obtained for n = 247 participants (i.e. two blood samples). For the analysis, mothers with the most severe depressive symptoms were selected, based on the longitudinal EPDS scores (23.53% with mild symptoms, 47.06% with moderate symptoms, and 29.42% with severe symptoms; (23). The PPD diagnosis was made in the clinical interview based on the DSM-5 criteria (n = 17; as well as a euthymic control group of mothers with stable low EPDS scores (n = 24).

### DNA Isolation from whole human blood

For the isolation of genomic DNA, 300 µL of whole human blood was thawed and suspended in 700 µL of RIPA lysis buffer (150 mM NaCl, 50 mM Tris/HCl; pH 8.0, 1% IGEPAL CA-630, 10% sodium deoxycholate, 10% SDS). The samples were then substituted with 5 µL proteinase K (Sigma-Aldrich) and incubated overnight at 52°C and 1000 rpm in the ThermoMixer (Eppendorf). DNA extraction was performed by adding 1000 µL of phenol-chloroform isoamyl alcohol (25:24:1, Sigma-Aldrich) to the lysate. The mixture was incubated for five minutes at room temperature first and then centrifuged at 15,000 rpm for 15 min. Subsequently, the resulting DNA-containing aqueous phase was transferred to a fresh Eppendorf tube, and an equal volume of 100% ethanol was added. Finally, the DNA was purified using the PureLink™ Genomic DNA Mini Kit (Invitrogen) according to the manufacturer’s protocol.

Genome-wide DNA methylation analysis was performed using the Infinium Human Methylation BeadChip (Illumina), following the manufacturer’s standard workflow. Briefly, 500 ng of genomic DNA per sample underwent bisulfite conversion using the Zymo EZ-96 DNA Methylation Kit (Zymo Research, Irvine, CA, USA). After conversion, DNA was amplified, fragmented, and purified before hybridization to the BeadChip arrays. The arrays were then washed and scanned on the Illumina iScan System. All array processing was conducted at the Core Facility Genomics, Helmholtz Munich.

### Analysis of Differential DNA Methylation

Illumina Infinium MethylationEPIC v2 BeadChip data were analyzed using the ChAMP package (v2.x) in R. Raw β-values and sample metadata were imported using champ.load(), with filtering for probes on X/Y chromosomes disabled. Quality control was performed using champ.QC() to assess sample quality and β-value distributions. Normalization was then applied via champ.norm() using the BMIQ method to correct for probe-type bias. Batch effect assessment was conducted using singular value decomposition (champ.SVD()).

Differential methylation analysis between conditions (postpartum depression vs. healthy controls at timepoints t0 and t4) was performed using champ.DMP(), which employs a limma-based linear model approach. CpG sites with a Benjamini-Hochberg adjusted p-value ≤ 0.05 and |Δβ| ≥ 0.10 were considered differentially methylated. For comparisons yielding no significant results at this threshold, the adjusted p-value cutoff was relaxed to 1.0 to capture all nominally differential sites.

### Genomic Annotation and Regulatory Context

All genomic annotations were mapped to the human reference genome GRCh38/hg38. Probe coordinates were obtained from the IlluminaHumanMethylationEPICv2anno.20a1.hg38 annotation package. DMP CpG IDs were matched to manifest coordinates either by probe identifier or, when necessary, by genomic position using GenomicRanges::findOverlaps().

CpG island context was defined using UCSC cpgIslandExt annotations: CpG islands were obtained directly from the UCSC table; shores were defined as regions within 2 kb flanking islands; shelves as regions 2–4 kb from islands; and all remaining regions classified as open sea.

Regulatory element annotation employed a hierarchical classification scheme using TxDb.Hsapiens.UCSC.hg38.knownGene and ENCODE Registry of cis-Regulatory Elements (cCREs). Promoters were defined as regions spanning −2,000 to +200 bp relative to transcription start sites. Enhancers were identified from ENCODE SCREEN cCRE annotations, specifically selecting enhancer-like signatures (pELS and dELS). Gene bodies encompassed the full transcribed region from TSS to transcription end site. CpGs were assigned to regulatory classes hierarchically: promoter, then enhancer, then gene body, with remaining sites classified as intergenic. Overlaps between CpG coordinates and regulatory features were determined using GenomicRanges::findOverlaps() with strand information ignored.

Gene proximity information was calculated using GenomicRanges::nearest() to identify the closest gene (by Entrez ID) for each CpG, with gene symbols retrieved from org.Hs.eg.db. Distance to the nearest transcription start site was computed for all probes.

Genomic annotations were performed using the annotatr package for standardized feature retrieval, with additional custom processing via GenomicRanges, GenomicFeatures, and rtracklayer packages. Probes lacking valid genomic coordinates were excluded from annotation analyses.

HCC Analysen: The average hair growth is considered to be 1 cm/month, with the posterior vertex region of the head showing the least variation in growth rates (24). Therefore, to reflect <u>cortisol</u> exposure over the last trimester of pregnancy and the three months postpartum , the first 3 cm of the hair segment was taken immediately following childbirth and after 12 weeks postpartum. The hair samples were stored in aluminum foil to prevent further contamination and were analyzed by means of the automatized online SPE LC–MS Method (25) at the Institute of Occupational Medicine of the University Hospital RWTH Aachen. In brief, approximately 50 mg of the hair was minced and washed with <u>isopropanol</u>, extracted with a 4-fold deuterium isotope-labeled internal standard of cortisol and methanol for 24 h and then analyzed with liquid chromatography triple quadrupole mass spectrometry using an ion trap (Agilent Technologies 1200 infinity series—QTRAP 5500 ABSciex, Darmstadt, Germany). A detailed description of the chemical processes during the analyses with SPE LC-MS^3^ and method validation is provided in (25). The limits of quantification were 0.05 ng/ml or 2 pg/mg hair, respectively.

## Results

### Group Characteristics and Longitudinal Stress Profiles in PPD vs. Controls

Out of 827 initially recruited postpartum women, a complete longitudinal dataset including two blood samples was available for 247 participants. From this cohort, we identified 17 mothers with significant depressive symptoms and a confirmed diagnosis of PPD, and 24 mothers with stable euthymic profiles over the 12-week period. The clinical status was determined based on the Edinburgh Postnatal Depression Scale (EPDS) scores (Figure 1d). We investigated group differences in subjectively perceived stress. Over the course of 12 weeks, participants with PPD had higher subjectively perceived stress experience compared to controls across all time points (Figure 1b; GEE Wald test: Group p=6.359×10^⁻17^), as well as a consistently lower Mood score (Figure 1a; GEE Wald test Group p= 1.077x10^-15^), indicating sustained stress exposure and symptom burden throughout the postpartum period.

**Figure 1:**
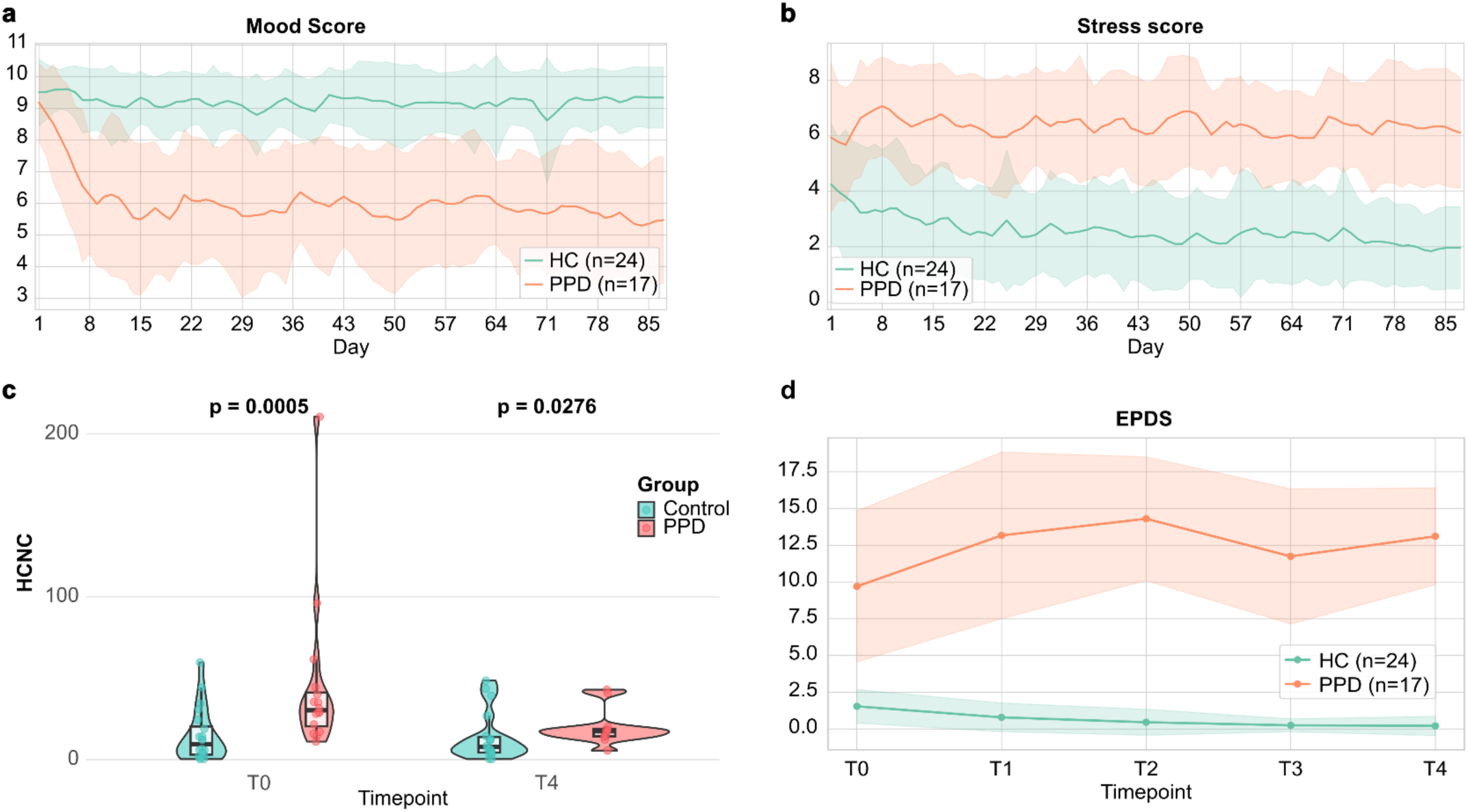
Clinical and biological parameters of patients with postpartum depression and healthy controls. **(a)** Self-reported mood scores from 1 to 10 throughout the study. **(b)** Perceived stress scores assessed throughout the study for control and PPD groups (same colour coding as panel a). The PPD group exhibits consistently higher stress scores than controls across all postpartum time points. **(c)** Hair cortisone concentration (HCNC) at T0 and T4, stratified by group (control: teal, PPD: orange). Individual data points are shown. Significant differences between-group differences are present at both time points (T0: p = 0.0005; T4: p = 0.0276), with PPD patients exhibiting persistently elevated cortisone levels. **(d)** Edinburgh Postnatal Depression Scale (EPDS) scores across time points T0–T4 for the control (teal, n = 24) and PPD (orange, n = 17) groups. Lines represent group means; ribbons indicate standard deviation. The PPD group maintains persistently elevated scores throughout the study period, whereas controls remain near zero.

Descriptive cohort characteristics are summarized in Table 1. The average age was comparable between the groups, with HCs averaging 33.46 years (SD = 4.33) and PPD participants 32.18 years (SD = 6.24). Gestational age at delivery was similar across groups (HC: 38.83 weeks, SD = 1.37; PPD: 38.65 weeks, SD = 1.54), as was mean birth weight of the newborns (HC: 3364, SD = 508.32; PPD: 3235, SD = 444.86). With respect to birth mode, spontaneous vaginal delivery was more frequent among controls (66.67%) than in the PPD group (41.18%), whereas emergency C-sections occurred more frequently in the PPD group (17.65%; HC: 4.17%). Elective C-sections were reported with equal frequency in both groups (25% in HC; 29.41% in PPD), and ventouse-assisted deliveries were relatively rare (Table 1).

**Table 1:** Descriptive statistics of the study cohort, shown for the whole sample (N = 37), healthy controls (HC, n = 19), and patients with postpartum depression (PPD, n = 18). Variables include maternal age, gestational age, child’s birth weight, birth mode, breastfeeding intention at T0 and breastfeeding status at T4, number of children, and relevant obstetric characteristics; values are reported as means with standard deviation or percentages as appropriate, with p values indicating between-group differences.

Breastfeeding intention at baseline (T0) was high in both groups (HC: 83.33%; PPD: 94.12%), and breastfeeding continuation at 12 weeks postpartum remained high (HC: 79.17%; PPD: 82.35%), suggesting broadly comparable early caregiving context (Table 1).

Parity and sociodemographic factors were likewise similar: the average number of children was nearly identical (HC: 1.63; PPD: 1.77) and approximately half of each group were first-time mothers (58.33% of HCs; 47.06% of PPDs; Table 1).

Educational attainment and marital status did not differ substantially between the groups, although a somewhat higher proportion of PPD participants reported secondary education being the highest level (Table 1).

In contrast, pronounced differences emerged in psychiatric history (p < .001): 95.83% of controls reported no prior psychiatric diagnosis, compared to only 47.06% of the PPD group. In the PPD cohort, 47.06% had a history of depression and 5.88% reported prior anxiety disorders. Family psychiatric history was more prevalent among women with PPD (47.06%) than in controls (12.5%; p = .014), consistent with known risk gradients for PPD. Together, these data confirm that the selected PPD and HC groups are broadly comparable in obstetric and sociodemographic characteristics but differ strongly in psychiatric vulnerability and postpartum stress trajectories.

### Widespread postpartum DNA methylation remodeling in healthy mothers versus attenuated changes in PPD

We first assessed longitudinal global blood methylome changes across the postpartum period between T0 (2-3 days postpartum) and T4 (12 weeks postpartum) separately in healthy controls and PPD patients. Genome-wide DNA methylation was profiled in peripheral blood of healthy mothers (HC, n=24) and women diagnosed with postpartum depression (PPD, n=17) using the Illumina Infinium MethylationEPIC v2 BeadChip. After quality control, normalization (BMIQ), and batch assessment with the ChAMP pipeline in R, differential methylation analysis was performed using limma-based models (champ.DMP) with Benjamini-Hochberg adjustment.

Healthy postpartum women exhibited widespread hypermethylation from T0 to T4, with thousands of CpG sites meeting the predefined thresholds (adjusted p ≤ 0.05 and |Δβ| ≥ 0.10; Fig. 2a). This is also reflected by the strong deviation of QQ-plots from the expected null distribution (Supplemental Fig. 1a). In total, 16,805 CpG sites surpassed the 10% methylation-change threshold and became hypermethylated, whereas 169 sites showed hypomethylation (Fig. 2a, left panel). In contrast, women with PPD showed markedly fewer significant DMPs with 10,436 hypermethylated and 120 hypomethylated sites identified between T0 and T4 (Fig. 2a). Their QQ-plots (Supplemental Fig. 1a, red) follow closer to the null distribution compared to the Control. This indicates a blunted epigenetic response to the postpartum transition. The large majority of the hypermethylated sites in PPD (10,410 sites) were also hypermethylated in HCs, representing the core temporal methylation response common to all postpartum women. However, 6,391 hypermethylated DMPs were unique to healthy controls, while only 362 were specific to the PPD group (Fig. 2a, right panel), suggesting that PPD may involve attenuated or dysregulated methylation responses. For the hypomethylated sites, 72 sites were found in both groups, whereas 99 sites were only detected in HCs and 63 sites in PPD (Fig. 2a, left panel).

**Figure 2:**
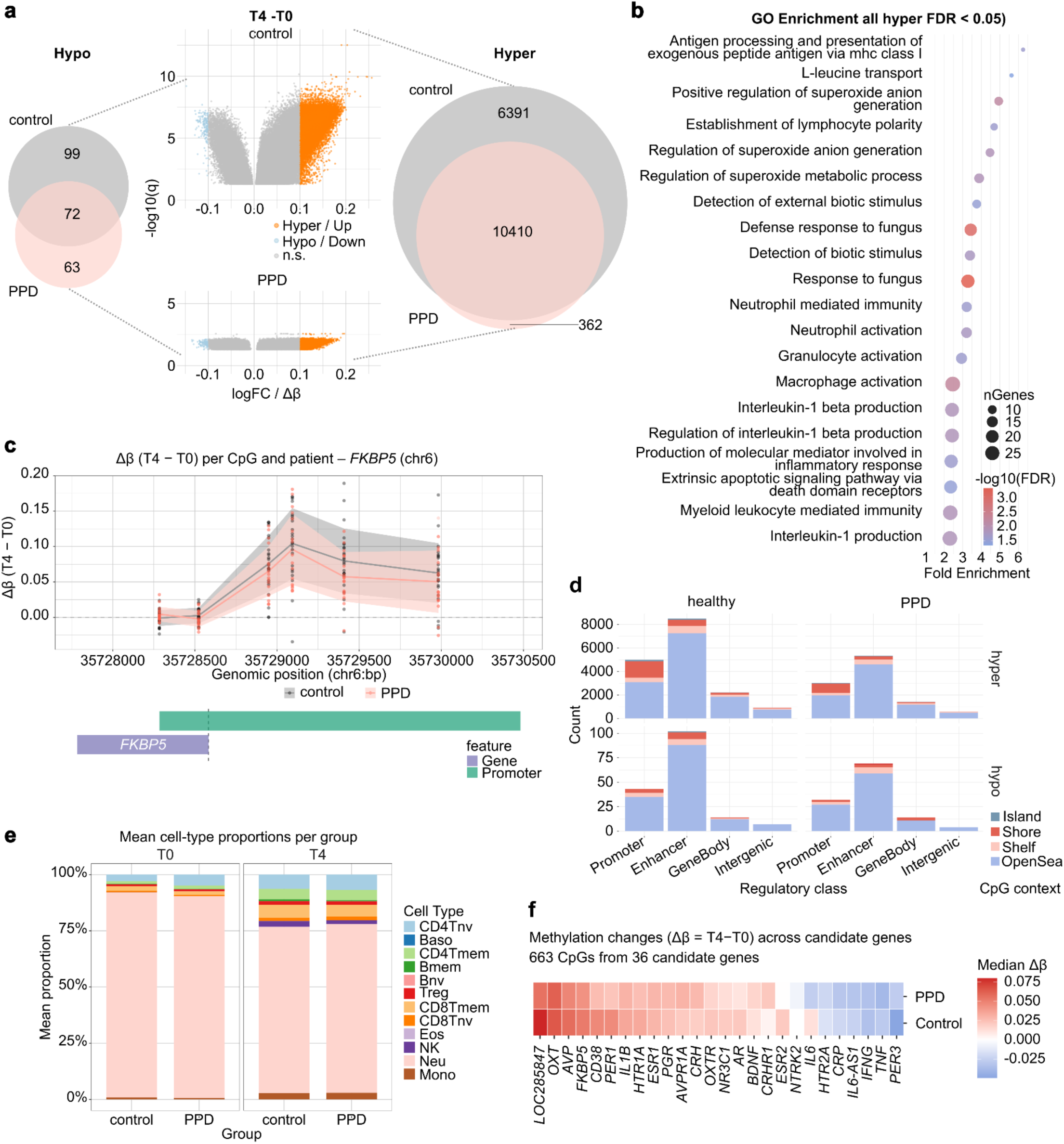
Longitudinal blood DNA methylation profiling in PPD patients and controls using the Infinium MethylationEPIC array. **(a)** Volcano plots of longitudinal within-group methylation change (Δβ = T4 − T0) for controls (top) and PPD patients (bottom). The x-axis shows Δβ, the y-axis −log10(q); hypermethylated CpGs (Δβ > 0) are orange, hypomethylated (Δβ < 0) blue, non-significant grey; dashed lines mark significance thresholds. Venn diagrams (right) summarize significantly changing CpGs per group (q < 0.05, |Δβ| > 0.1), with 6,391 sites unique to controls, 362 unique to PPD, and 10,410 shared. **(b)** Gene Ontology enrichment for hypermethylated CpGs pooled across groups (FDR < 0.05). Bar length indicates fold enrichment, colour encodes −log10(FDR), and point size the number of associated genes; enriched terms are dominated by immune and inflammatory processes (for example interleukin-1 production, neutrophil and granulocyte activation, myeloid leukocyte-mediated immunity). **(c)** Longitudinal methylation change (Δβ = T4 − T0) at the FKBP5 locus (chr6: ∼35,728,000–35,730,500), shown per CpG and patient. Trajectories for controls (teal) and PPD (red/pink) are overlaid with group means ± SD; gene structure and promoter annotation appear below the x-axis. **(d)** Numbers of DMPs by regulatory class (promoter, enhancer, gene body, intergenic) in controls and PPD, subdivided by CpG island context (island, shore, shelf, open sea) and direction of change (hyper/hypo). **(e)** Mean blood cell-type proportions at T0 and T4 for control and PPD groups, estimated by reference-based deconvolution. Each stacked bar shows the mean proportion of 12 cell types (CD4Tnv, Baso, CD4Tmem, Bmem, Bnv, Treg, CD8Tmem, CD8Tnv, Eos, NK, Neu, Mono); monocytes (Mono, pink) dominate in all groups, with no marked shifts over time. **(f)** Heatmap of Δβ (T4 − T0) for 663 CpGs in 36 literature-derived PPD candidate genes, shown for PPD (top) and controls (bottom). Colour encodes median gene-wise Δβ on a shared scale (−0.025 to 0.075); genes with the largest positive Δβ in PPD include AVP and FKBP5.

**Figure 3:**
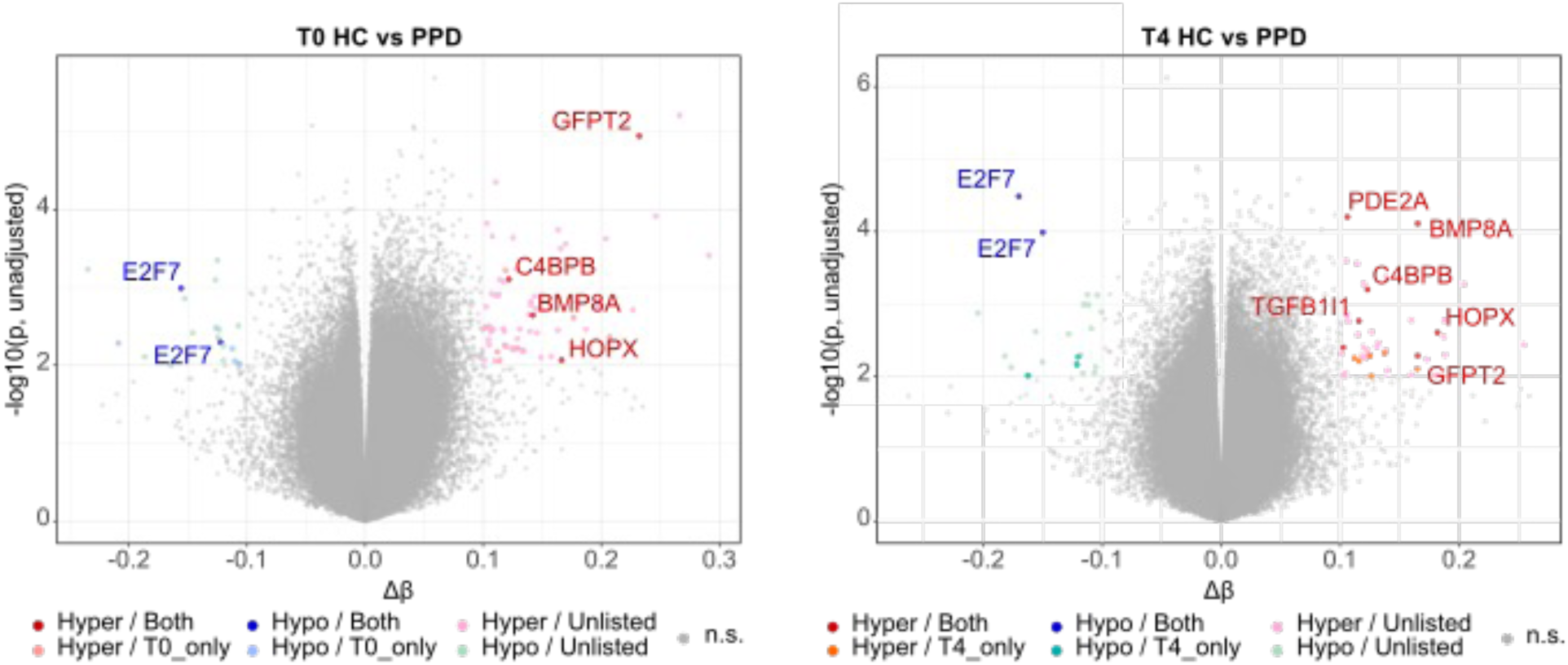
Unadjusted differential DNA methylation between healthy controls and PPD patients Volcano plots at T0 (left panel) and T4 (right panel) comparing beta values between healthy controls (HC) and PPD groups using unadjusted p values. Sites with p < 0.01 and |Δβ| > 0.1 are coloured according to whether they are dysregulated at one or both time points; sites dysregulated at both time points and mapping to a gene are annotated.

Because DNA methylation profiles obtained from whole blood can be strongly influenced by changes in cell-type composition, we next examined whether the observed longitudinal methylation shifts might reflect underlying immune remodeling. We applied EpiDISH-based cell-type deconvolution using the Cent12CT.m reference panel to estimate the relative abundance of major immune cell populations at both timepoints (T0 and T4) in healthy and PPD participants (Figure 2e). As expected for peripheral blood samples, both cohorts displayed comparable immune cell compositions at each time point. Across time, however, both healthy and PPD groups exhibited pronounced postpartum shifts in immune cell proportions, with a clear increase in CD4⁺ and CD8⁺ T-cell subsets, including both naïve and memory populations, from T0 to T4. These changes are in line with prior reports of postpartum immune recovery following pregnancy-induced immunomodulation, suggesting that part of the longitudinal methylation remodeling captured in whole blood may be driven or amplified by physiological immune reconstitution (5). Because the immune cell compositions were highly similar between the groups at both time points, the more pronounced hypermethylation signatures in healthy women from T0 to T4 cannot be attributed to gross differences in blood cell makeup.

To better understand the biological processes associated with these postpartum methylation trajectories, we performed Gene Ontology (GO) enrichment analyses on genes acquiring hypermethylated DMPs at T4 compared to T0. In the combined sample, hypermethylated genes were strongly enriched for immune-related terms, including neutrophil activation, phagocytosis, myeloid leukocyte activation, and regulation of adaptive immune responses (Figure 2b). DMPs unique to healthy controls showed enrichment for overlapping immune terms such as neutrophil degranulation, leukocyte activation, and inflammatory responses (Supplementary Figure 1b). Hypermethylated DMPs overlapping between PPD and control groups similarly converged on immune and inflammatory processes, with the strongest enrichments (fold > 3, FDR < 0.05) for positive regulation of superoxide anion production, defense responses to fungi and bacteria, neutrophil/granulocyte chemotaxis, degranulation and activation, phagocytosis/endocytosis, myeloid leukocyte differentiation, and adaptive immunity (Supplementary Figure 1c). Together with the cell-composition estimates, these findings support the view that postpartum blood methylation trajectories predominantly capture an orchestrated adjustment of innate and adaptive immune programs, which appears robust and coordinated in healthy women but attenuated and more heterogeneous in those who develop PPD. Genomic context analysis revealed that nearly 30% of hypermethylated DMPs localized to promoter regions (defined as 2 kb upstream to 200 bp downstream of the transcription start site), with very similar proportions in healthy controls and PPD samples (Figure 2d). As a representative example, the FKBP5 gene on chromosome 6 showed a pronounced gain of methylation in its promoter region across multiple CpG sites in both groups, with individual patient trajectories illustrating the consistency of this change over time (Figure 2c). More than 50% of hypermethylated DMPs were assigned to enhancer regions, approximately 14% to gene bodies, and about 5% to intergenic regions in both cohorts (Figure 2d). The genomic context of hypomethylated sites followed a similar pattern, without gross differences between groups, and the chromosomal distribution of DMPs was likewise comparable (Supplementary Figure 1d).

The reduced number of significant DMPs in the PPD group, together with the generally lower –log₁₀(adjusted p) values in the PPD volcano plots, pointed toward increased inter-individual variability in methylation dynamics. This suggests that women with PPD undergo a less coordinated epigenetic response to the postpartum transition compared to healthy controls.

To quantify this effect, we focused on CpG sites that were significantly altered over time in healthy controls and compared the variance of methylation changes between groups. For all CpG sites classified as hyper- or hypomethylated in controls, we computed the variance of methylation values separately for HC and PPD groups at both timepoints (T0 and T4). Relative variability was defined as the difference in variance (VarPPD − VarHC), such that positive values indicate higher variability in PPD and negative values higher variability in controls (Supplemental Figure 2a).

Among sites that became differentially methylated in controls from T0 to T4, the majority (15,012 CpGs) exhibited greater inter-individual variance in PPD patients already at the first timepoint (T0), whereas only 1,960 sites showed higher variance in the HC samples. A similar imbalance was observed at T4: 15,683 CpGs displayed higher variance in PPD compared to control samples, with only 1,289 sites showing lower variance in PPD. We repeated this analysis restricting to the 6,391 sites that were significantly hypermethylated from T0 to T4 in HCs but not in the PPD cohort (Supplemental Figure 2c). Again, more than 5,000 sites showed higher variation in PPD compared to HC samples at both T0 and T4.

### Temporally stable candidate blood methylation markers with potential predictive value for PPD

Applying rigorous multiple testing correction across the entire array, none of the individual CpG sites was found to reach genome-wide significance in our relatively small sample of 41 women, which is fully in line with many previous epigenome-wide studies of depression and postpartum depression using comparable cohort sizes (26–28). Given the modest effect sizes typically observed for peripheral DNA methylation markers and the very large number of tests, such studies are often underpowered to detect single CpGs at stringent FDR thresholds. Therefore, they rely largely on complementary, biology-informed analytical strategies, including the use of nominally associated CpGs, region-based tests (differentially methylated regions), gene- and pathway-level aggregation, and overlap with prior GWAS or candidate-gene findings (29).

In line with the current practice in epigenome-wide studies of (postpartum) depression, which frequently lack genome-wide significant single-CpG hits in modestly sized cohorts and therefore rely on nominally associated sites, region-based tests and gene- or pathway-level aggregation (26,30), we focused on CpG sites that showed consistent direction and magnitude of methylation differences between PPD cases and controls at both T0 and T4. We also examined whether the corresponding genes cluster in biologically meaningful pathways and include the loci previously implicated in depression and stress-related psychopathology.

In our cross-sectional analysis of CpGs that were consistently hypermethylated in women with PPD at both T0 and T4, we identified a set of 57 loci mapping to protein-coding genes and non-coding transcripts with functions in synaptic signaling, neurodevelopment, epigenetic regulation and stress/inflammatory pathways. Several of these genes have been implicated previously in major depressive disorder or stress-related psychopathology, whereas others represent biologically plausible but hitherto unreported candidates in the context of postpartum depression(26,28,31).

Notably, our “T0+T4” gene set includes regulators of glutamatergic neurotransmission and cAMP/cGMP signaling such as GRIN3A, GRIK2 , TACR3 and PDE2A (32–34). GRIN3A encodes the GluN3A NMDA receptor subunit, and experimental disruption of GluN3A has been shown to induce depression-like behavior in rodents, with restoration of GluN3A levels ameliorating affective phenotypes (35). GRIK2 and related glutamatergic genes have been associated with specific depressive symptom dimensions and antidepressant treatment response, while pharmacological inhibition of PDE2A exerts robust antidepressant- and anxiolytic-like effects in animal models (36). These findings are consistent with the view that subtle epigenetic alterations in glutamatergic and second-messenger signaling pathways contribute more broadly to vulnerability for depressive phenotypes (26,31).

Among intracellular signaling modulators, we observed persistent hypermethylation at an upstream CpG in RGS4, a key regulator of G-protein-coupled receptor signaling that has repeatedly been linked to major depression, schizophrenia and altered prefrontal function (37–39). Experimental manipulation of RGS4 in mouse prefrontal cortex modulates stress-related behavior and ketamine-sensitive depression-like phenotypes, highlighting RGS4 as a mechanistically relevant node at the intersection of monoaminergic, glutamatergic and stress pathways. In the context of PPD, RGS4 hypermethylation at both T0 and T4 may thus reflect a pre-existing vulnerability factor or a stable epigenetic adaptation within the prefrontal circuits.

In addition to these previously implicated depression-related genes, our persistent hypermethylation signature contains multiple transcription factors and developmental regulators (e.g. DMRTA1, NR2F2, CRX, HOPX, ZNF311) as well as components of TGF-β/BMP and lipid-mediator signaling (BMP8A, TGFB1I1, ALOX15, PLA2G12A). While these specific loci have not been systematically studied in PPD, TGF-β/BMP signaling, neurodevelopmental transcriptional programs and inflammatory/lipid pathways are consistently highlighted in epigenetic and transcriptomic studies of depression and stress exposure, suggesting that our findings extend these themes into the postpartum period. Finally, several genes in the T0+T4 set encode epigenetic and post-transcriptional regulators (YTHDF2, METTL22, CSTF2 and multiple lncRNAs), in line with emerging evidence that RNA modifications and non-coding RNAs contribute to depression-related phenotypes. Together, these observations indicate that the subset of CpGs hypermethylated at both time points captures a biologically coherent core of pathways with prior links to depression and stress, while also nominating novel PPD candidate genes within these pathways that warrant targeted validation in larger and independent cohorts.

## 3. Discussion

In this longitudinal epigenome-wide study of postpartum women, we show that healthy mothers undergo a robust and coordinated wave of DNA methylation remodeling over the first 12 weeks after delivery, whereas women who develop postpartum depression (PPD) display an attenuated and more heterogeneous epigenetic response. This divergence is not explained by sociodemographic or obstetric differences, nor by gross changes in blood cell composition, but instead suggests a dysregulated epigenetic adaptation of immune-related pathways in PPD, accompanied by increased inter-individual variability and subtle, temporally stable alterations in genes such as GFPT2, GRIN3A, GRIK2, TACR3, PDE2A and RGS4.

These findings indicate that women with PPD exhibit a markedly less coordinated epigenetic response to the postpartum period. While healthy mothers show a robust and relatively uniform hypermethylation trajectory, PPD mothers display highly heterogeneous methylation profiles, already evident at delivery and persisting into early postpartum weeks. Such elevated inter-individual variability may reflect dysregulated physiological adaptation, including altered stress reactivity, immune activation, or hormonal signaling, which likely disrupt the normal consolidation of postpartum epigenetic states.

### Impaired coordination of postpartum immune-epigenetic adaptation in PPD

Healthy women showed widespread hypermethylation from 2-3 days to 12 weeks postpartum, with thousands of CpGs surpassing stringent thresholds and enriched for innate and adaptive immune processes, consistent with the view of the postpartum period as a phase of intense immunological recalibration. Cell-type deconvolution revealed the expected shift from a neutrophil-dominated profile toward higher CD4⁺/CD8⁺ T-cell proportions in both groups, indicating that broad immune-related hypermethylation partly reflects physiological immune reconstitution. Despite similar estimated immune cell compositions at each time point, CpGs that remodeled robustly in controls showed substantially higher inter-individual variability in PPD at both T0 and T4, especially for sites hypermethylated over time only in healthy women. These findings support a model in which healthy mothers mount a relatively synchronized immune-epigenetic response to the postpartum transition, whereas women with PPD show a blunted and poorly coordinated response, in line with models of impaired allostatic regulation of immune-inflammatory pathways in depression (5,28,40–42).

### Genomic context and convergence with stress-related genes

Nearly one-third of hypermethylated DMPs were located in promoters and more than half in enhancers, with similar distributions in both groups, indicating that PPD is not defined by a distinct methylation architecture but by quantitative and coordination differences in a largely shared immune-epigenetic program. FKBP5, a key regulator of glucocorticoid receptor sensitivity, exemplified this pattern by showing increased promoter methylation in both healthy and PPD mothers, consistent with prior links between FKBP5 methylation, HPA-axis dysregulation and stress-related psychopathology (20,26,43).

### Temporally stable candidate blood methylation markers with potential relevance for PPD

No individual CpG reached epigenome-wide significance in direct PPD versus control comparisons at either time point. This was expected given the modest sample size and high dimensionality, mirroring previous difficulties in identifying genome-wide significant single-CpG hits in PPD. We therefore focused on CpG sites that showed consistent direction and magnitude of methylation differences between PPD cases and controls at both T0 and T4, and examined whether the corresponding genes cluster in biologically meaningful pathways with prior links to depression and stress. Using this approach, we identified 57 CpGs that were nominally hypermethylated in PPD at both time points and mapped to genes involved in synaptic signaling, neurodevelopment, epigenetic regulation and stress/inflammatory pathways, including GRIN3A, GRIK2, TACR3, PDE2A and RGS4. Experimental and genetic evidence implicates these loci in glutamatergic neurotransmission, cAMP/cGMP signaling and GPCR modulation, and links RGS4 specifically to depression-related behavior and prefrontal function, suggesting that subtle, temporally stable epigenetic alterations in these pathways may contribute to PPD vulnerability (28,31–39,44).

Beyond these previously implicated depression-related genes, the stable hypermethylation signature includes transcription factors and developmental regulators (e.g. DMRTA1, NR2F2, CRX, HOPX, ZNF311), components of TGF-β/BMP and lipid-mediator signaling (BMP8A, TGFB1I1, ALOX15, PLA2G12A), and epigenetic/post-transcriptional regulators (YTHDF2, METTL22, CSTF2 and several lncRNAs), extending themes from broader depression and stress-related epigenetic studies into the postpartum period. Given the limited power and lack of FDR-significant group differences, these candidates remain hypothesis-generating, but the convergence on glutamatergic signaling, GPCR/second-messenger modulation, developmental transcription factors and immune-related regulators suggests a biologically coherent core of pathways for targeted replication and integration into multi-marker prediction models alongside established PPD methylation markers such as TTC9B and HP1BP3 (26–28,31).

### Comparison with previous PPD methylation studies

Previous blood-based PPD methylation studies, often focusing on antenatal prediction, have highlighted TTC9B, HP1BP3, NR3C1, OXTR and immune and stress-related pathways, and have achieved only moderate predictive accuracy. Our study differs by characterizing the postnatal trajectory from delivery to 12 weeks postpartum, quantifying inter-individual variability in methylation dynamics and integrating immune cell deconvolution, but converges conceptually in emphasizing immune and inflammatory programs as central to PPD biology. By nominating GFPT2, GRIN3A, GRIK2, TACR3, PDE2A and RGS4 as consistently altered across time in PPD, our work extends candidate pathways beyond estrogen-sensitive markers and suggests that metabolic-epigenetic coupling and nuclear lncRNA function may also be relevant. Likewise, transcriptome-wide studies in large peripartum cohorts report immune dysregulation as a core correlate of peripartum depressive symptoms. These observations, together with our findings of blunted and less coordinated immune-related methylation remodeling in PPD, converge on immune dysregulation as a central feature across molecular layers (42,45).

### Limitations

Our candidate-based analyses should be viewed as hypothesis-generating, requiring replication, given that the modest PPD sample size (n = 17) limits power for epigenome-wide association. It also increases the risk of false negatives and positives, and likely contributes to the lack of epigenome-wide significant DMPs. Whole-blood measurements and deconvolution cannot fully separate within-cell-type regulatory changes from subtle shifts in cell composition or activation state. To refine cellular resolution, future research would need to use sorted cells, single-cell methylation or parallel transcriptomics. The two-time point design captures longitudinal change and temporal stability for selected genes, but not detailed dynamics or non-linear trajectories. In addition, the observational nature of the study precludes causal inference regarding methylation changes, immune adaptation and depressive symptoms. Finally, the cohort stems from a specific cultural and healthcare context, which may limit generalizability, underscoring the need for replication in diverse populations (5,27,43).

### Conclusions and outlook

PPD appears to be associated less with a qualitatively distinct methylation signature than with a failure to mount a robust and coordinated immune-epigenetic adaptation to the postpartum period. Healthy mothers show a synchronized hypermethylation trajectory in immune and stress-related genes that parallels postpartum immune recovery, whereas women with PPD exhibit attenuated and highly variable changes, with temporally stable hypermethylation at genes such as GFPT2, GRIN3A, GRIK2, TACR3, PDE2A and RGS4 that may form part of a metabolic-epigenetic axis contributing to postpartum mood vulnerability. Larger samples, higher temporal resolution and cell-type-specific analyses will be critical to validate these candidates, integrate them with established PPD biomarkers and clarify whether aspects of postpartum immune-epigenetic adaptation can be leveraged for risk stratification or preventive intervention in at-risk mothers (27,46).

## Acknowledgements

We acknowledge the technical support of Core Facility Genomics at Helmholtz Munich. We gratefully acknowledge Jana Egner-Walter and Magdalena Vogl for purifying genomic DNA from blood samples as part of their HIWI positions, and Mira Jakovcevski for initially training Magda in the standardized protocol used in this study. We further thank the Biomaterial Bank of the University Hospital Aachen, located at the Institute for Pathology, for storing the blood samples. We sincerely thank all women who participated in this study for their invaluable contribution.

This research was funded by the Deutsche Forschungsgemeinschaft (DFG, German Research Foundation)- 368482240/GRK2416, ZI 1224/19-1, dedicated to G.Z.B., and Grant numbers: 410314797, 512021469) dedicated to N.C.. The study was further funded under the Excellence Strategy of the Federal Government and the Länder (SFASIA002) dedicated to G.Z.B..

The Authors declare no conflict of interest.

