## Supplementary Figures for "Longitudinal blood DNA methylation profiling reveals disrupted immune–epigenetic adaptation and candidate stress-related loci in postpartum depression"

### Supplementary Figure 1

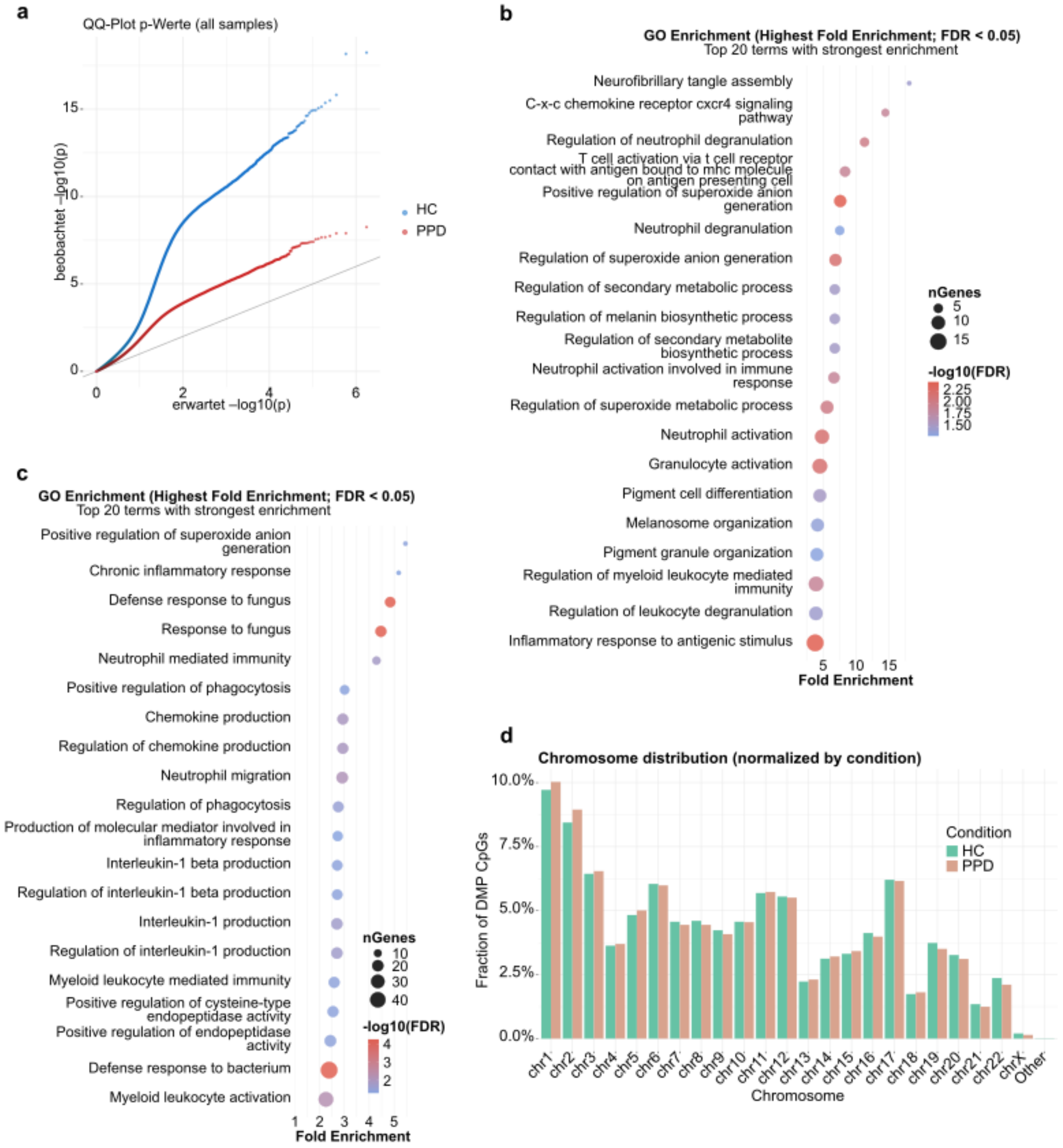

**Supplementary Figure 1**: **EPIC array–based blood DNA methylation QQ plots, GO enrichment, and chromosomal distribution at T0 and T4 in postpartum depression and control groups.**

**(a)** QQ plot of observed vs expected −log10(p) values for all EPIC CpGs, shown separately for healthy control (teal) and PPD (orange) groups. Both deviate modestly from the diagonal at extreme values, indicating genuine biological signal without substantial p-value inflation.

**(b)** Top 20 GO terms by fold enrichment (FDR < 0.05) for CpGs significantly hypermethylated exclusively in controls (T4 vs T0; n = 6,391 control-only CpGs). Enriched terms include neurofibrillary tangle assembly, C–X–C chemokine receptor CXCR4 signalling, neutrophil degranulation, T-cell activation, melanosome and pigment granule organisation, and regulation of myeloid leukocyte-mediated immunity.

**(c)** Top 20 GO terms by fold enrichment (FDR < 0.05) for CpGs hypermethylated in both controls and PPD patients (shared, n = 10,410 overlapping CpGs). Enriched terms include positive regulation of superoxide anion generation, chronic inflammatory response, defence response to fungus, neutrophil-mediated immunity, chemokine production, interleukin-1 beta production, and regulation of endopeptidase activity, reflecting biological processes common to both groups across the postpartum period.

**(d)** Fraction of significant DMP CpGs per chromosome, normalised within condition (control/PPD), showing broadly similar chromosomal distributions between groups with minor enrichments at specific chromosomes.

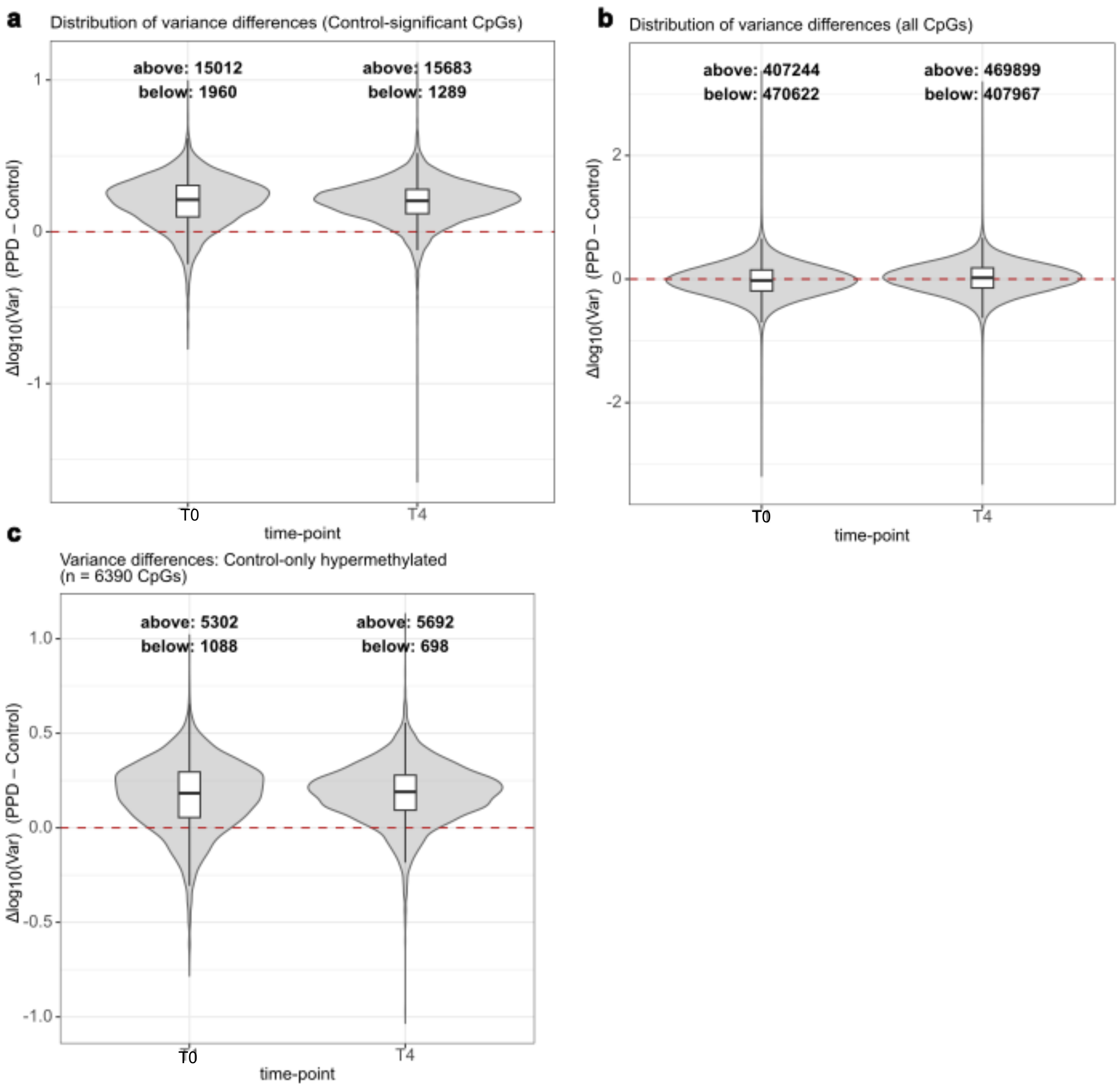


**Supplementary Figure 2: DNA methylation variance between postpartum depression patients and healthy controls**

**(a)** Violin plots of Δlog10(variance) (PPD minus control) at T0 and T4, restricted to CpGs significant in the control group. Numbers above and below zero indicate counts of CpGs with higher or lower variance in PPD, respectively; at both time points, most control-significant CpGs show greater variance in PPD patients.

**(b)** Same analysis as in panel a, but computed across all EPIC array CpGs (~450K). Distributions are centred near zero, indicating no global variance difference between groups.

**(c)** Variance analysis restricted to control-only hypermethylated CpGs (n = 6,390). These CpGs show a consistent positive skew at both T0 and T4, indicating that sites hypermethylated in healthy individuals are systematically more variable in PPD patients.
