## Supplementary material for "Longitudinal blood DNA methylation profiling reveals disrupted immune–epigenetic adaptation and candidate stress-related loci in postpartum depression": Table 1

### Descriptive Statistics

| | Whole Sample (N = 37) | | | HC (n =19) | | | PPD | | | *p* |
|---|---|---|---|---|---|---|---|---|---|---|
| | **Mean** | **SD** | **Percent** | **Mean** | **SD** | **Percent** | **Mean** | **SD** | **Percent** | |
| Age | 32.92 | 5.17 | | 33.46 | 4.33 | | 32.18 | 6.24 | | .471 |
| Gestational age in weeks | 38.76 | 1.43 | | 38.83 | 1.37 | | 38.65 | 1.54 | | .686 |
| Child's birth weight (g) | 3395.49 | 487.87 | | 3364.17 | 508.32 | | 3234.59 | 444.86 | | .403 |
| Birth mode | | | | | | | | | | .277 |
| Spontaneous | n = 23 | | 56.1 | n = 16 | | 66.67 | n = 7 | | 41.18 | |
| Ventouse | n = 3 | | 7.32 | n = 1 | | 4.17 | n = 2 | | 11.77 | |
| C-Section | n = 11 | | 26.83 | n = 6 | | 25 | n = 5 | | 29.41 | |
| Emergency C-Section | n = 4 | | 9.76 | n = 1 | | 4.17 | n = 3 | | 17.65 | |
| Intent to breastfeed at t0 (yes) | n = 36 | | 87.81 | n = 20 | | 83.33 | n = 16 | | 94.12 | .299 |
| Breastfeeding at t4 (yes) | n = 33 | | 80.49 | n = 19 | | 79.17 | n = 14 | | 82.35 | .800 |
| Number of children | 1.68 | 0.85 | | 1.63 | 0.88 | | 1.77 | 0.83 | | .610 |
| 1 | n = 22 | | 53.66 | n = 14 | | 58.33 | n = 8 | | 47.06 | |
| 2 | n = 11 | | 26.83 | n = 6 | | 25 | n = 5 | | 29.41 | |
| 3 | n = 7 | | 17.07 | n = 3 | | 12.5 | n = 4 | | 23.53 | |
| 4 | n = 1 | | 2.44 | n = 1 | | 4.17 | | | 0 | |
| Secondary education | | | | | | | | | | .904 |
| Lowest (< 9 years) | n = 2 | | 4.88 | n = 1 | | 4.17 | n = 1 | | 5.88 | |
| Middle (10 - 12 years) | n = 16 | | 39.02 | n = 10 | | 41.67 | n = 6 | | 35.29 | |
| Highest (> 13 years) | n = 23 | | 56.1 | n = 13 | | 54.17 | n = 10 | | 58.82 | |
| Married (yes) | n = 30 | | 73.17 | n = 18 | | 75 | n = 12 | | 70.59 | .753 |
| Psychiatric History | | | | | | | | | | <.001 |
| no | n = 31 | | 75.61 | n = 23 | | 95.83 | n = 8 | | 47.06 | |
| Depression | n = 8 | | 19.51 | | | 0 | n = 8 | | 47.06 | |
| Anxiety Disorder | n = 2 | | 4.88 | n = 1 | | 4.17 | n = 1 | | 5.88 | |
| Family Psychiatric History (yes) | n = 11 | | 26.89 | n = 3 | | 12.5 | n = 8 | | 47.06 | .014 |
